# Computer generation of fruit shapes from DNA sequence

**DOI:** 10.1101/2022.09.19.508595

**Authors:** M. Pérez-Enciso, C. Pons, A. Granell, S. Soler, B. Picó, A.J. Monforte, L.M. Zingaretti

## Abstract

The generation of realistic plant and animal images from marker information could be a main contribution of artificial intelligence to genetics and breeding. Since morphological traits are highly variable and highly heritable, this must be possible. However, a suitable algorithm has not been proposed yet. This paper is a proof of concept demonstrating the feasibility of this proposal using ‘decoders’, a class of deep learning architecture. We apply it to Cucurbitaceae, perhaps the family harboring the largest variability in fruit shape in the plant kingdom, and to tomato, a species with high morphological diversity also. We generate Cucurbitaceae shapes assuming a hypothetical, but plausible, evolutive path along observed fruit shapes of *C. melo*. In tomato, we used 353 images from 129 crosses between 25 maternal and 7 paternal lines for which genotype data were available. In both instances, a simple decoder was able to recover expected shapes with large accuracy. For the tomato pedigree, we also show that the algorithm can be trained to generate offspring images from their parents’ shapes, bypassing genotype information. Data and code are available at https://github.com/miguelperezenciso/dna2image.

## Introduction

Shape and color patterns of animals and plant fruits are not only aesthetic features, but they also convey essential information on animal welfare or fruit quality and can be critical for consumer appreciation. Besides, plant and animal appearance have played a major role ever since domestication and many breeds and plant varieties were created based on morphology. Even today, breeders’ associations can spend much time in defining the ‘racial standard’. Often, domestication and breeding have untapped a range of shapes that was not present in the wild. The variability in morphology and colors in the dog is amazing compared to that of its wild ancestor the wolf. In plants, domestic squashes and gourds exhibit an enormous diversity in shapes whereas its wild counterparts produce small, rounded fruits only (Xanthopoulou *et al*. 2019). Today, dairy bull catalogs, a business worth millions of euros worldwide, usually present a picture of the bull in addition to its genetic evaluation. Bull catalogs usually include information on a ‘global’ conformation score that is an important part of the genetic value of the bull, and an indication of longevity. In many vegetables breeding programs, experienced breeders rely on their ‘eye’ to quickly discard unpromising experimental crosses.

Shape is easily modified by artificial selection and, unsurprisingly, has received much attention from the genetics community and the breeding industry (Tanksley 2004; Monforte *et al*. 2014). Tomato is perhaps the best studied species from a morphological point of view; numerous quantitative trait loci (QTL) and some causative genes affecting shape have been identified (Monforte 2014; Snouffer et al. 2020). Cucurbitaceae in turn have been less well studied, yet they allegedly display the largest morphological variability in the plant kingdom (Paris 2001). For instance, a whole sequencing effort of the different *C. pepo* morphotypes did reveal numerous SNP differences but no clear clue on causative loci for shape (Xanthopoulou *et al*. 2019).

The statistical analysis of shape has a long history in Evolution, which has fostered most of the analysis tools available today (Zelditch *et al*. 2004; Claude 2008; Klingenberg 2010). Traditional morphometrics is based on the analysis of summary statistics such as length, width, ratios, and areas (Brewer *et al*. 2006). Modern morphometrics, in turn, is based on the concept of ‘landmarks’ (Zelditch *et al*. 2004). A landmark is an anatomical position that can be identified in all samples, e.g., the tip of the nose in cattle. In landmark-based geometric morphometrics, the spatial information is contained in landmark coordinates. Shapes can then be compared once a common reference scale is found. This can be done via Generalized ‘Procrustes’ Analysis (GPA, Gower 1975), which consists of finding an optimal superimposition of several shapes such that distances between them are minimized.

In breeding, morphology research has focused so far on detection of quantitative trait loci (QTL) of shape-derived statistics (e.g., Monforte et al. 2014). These QTL often explain only a percentage of observed variability. This is not unexpected; a large body of literature shows that significant loci identified from genomewide association studies (GWAS) explain but a small percentage of genetic variability in complex traits (Wood *et al*. 2014; Robinson *et al*.2017; Visscher *et al*. 2017). Therefore, GWAS is not optimum for prediction. An alternative is to use all markers for prediction of some of the shape metrics (Tong *et al*. 2022). Nevertheless, shape is highly dimensional, and the QTL or genomic prediction approaches restrict the list of potential candidate genes by focusing on single univariate statistics. In addition, these summary statistics do not allow reconstructing the original shape and hampers the prediction of global appearance changes induced by selection.

Here we approach this issue from a holistic, opposite angle. We propose to reproduce expected shapes and textures that would result from a given individual’s DNA sequence. To that end, we explore algorithms based on deep learning tools. Note that, in contrast to standard descriptors of shape, the goal here is prediction given new DNA information rather than QTL search. Breeding is mainly concerned with prediction of future offspring performance and this proposal aligns with this target. This novel theoretical framework can have an important impact in breeding.

This paper is a proof of concept that the proposed approach is feasible, at least in simplified scenarios. We use a class of deep architectures, called ‘decoders’, to reproduce the expected shapes given a linear vector of causative polymorphisms and random SNPs. First, we show how a trained decoder is able to generate simple geometric forms (2D and 3D ellipses) followed by more realistic applications in cucurbitas and tomato fruits. We end by showing that, provided shapes are inherited through an ‘additive’ mechanism, the algorithm can predict offspring shapes based on parents’ images, bypassing genotype information. More sophisticated algorithms would be needed if shapes are not inherited ‘additively’.

## Material and methods

### Generation of simple 2D and 3D images

We first performed a simple experiment using 2D ellipse and 3D ellipsoid shapes to verify that the proposed architecture is useful. An ellipse can be defined by the lengths of its horizontal (x) and vertical (y) axes, plus a third axis z for 3D shapes. We drew 2D ellipses with cv2.ellipse() function from OpenCV python package (Bradski 2000) randomly varying x and y axis lengths, that is, ellipses differed in shape, size, and orientation. Images were black and white of size 64 x 64 pixels. The decoder network (described below) was trained using an input vector containing x/y ratio and ellipse size, i.e., the two ‘causative loci’, and 100 random uniformly distributed variables. The 100 random numbers were aimed at representing noise from DNA information that is unrelated to the ‘phenotype’, i.e., the image containing the ellipse.

We generated 3D ellipsoids as three-dimensional binary arrays using pymrt package (Metere and Möller 2017), array size was 32 x 32 x 32. As in the previous example, images were predicted from x, y, and z axes lengths plus 100 random uncorrelated variables. For representation of the 3D shapes, ellipsoid projections were drawn using the marching cubes algorithm as implemented in skimage (van der Walt *et al*. 2014) and the plot_trisurf package. However, since these 3D plots were not too accurate, we also plotted the ellipsoid sections across the x, y, and z axes. Both observed and predicted shapes were plotted.

### Cucurbit shapes

*C. pepo* fruits can adopt an enormous diversity of shapes (Figure 1A). This variability appeared only after domestication, since all wild fruits are small and round (Paris 1986). According to (Paris 1989, 2001), *C. melo* shapes may have followed several evolutive pathways. One pathway would be wild gourd (akin to pumpkin shape) → scallop → acorn; a second pathway would be wild gourd → marrow → straightneck → zucchini → cocozelle (Figure 1B). See also Figure 17 in (Paris 1989). We extracted contours from the ‘contours.png’ file, based in (Paris 1989) and available in GitHub (https://github.com/miguelperezenciso/dna2image/blob/main/images/contours.png), using OpenCV library (Bradski 2000). Contours were centered and 500 pseudo-landmarks were obtained with the algorithm in Zingaretti et al. (2021). Next, contours were aligned with a generalized procrustes algorithm implemented in python package ‘procrustes’ (Meng *et al*. 2022) and images were resized to 64 x 64 pixels.

**Figure 1:**
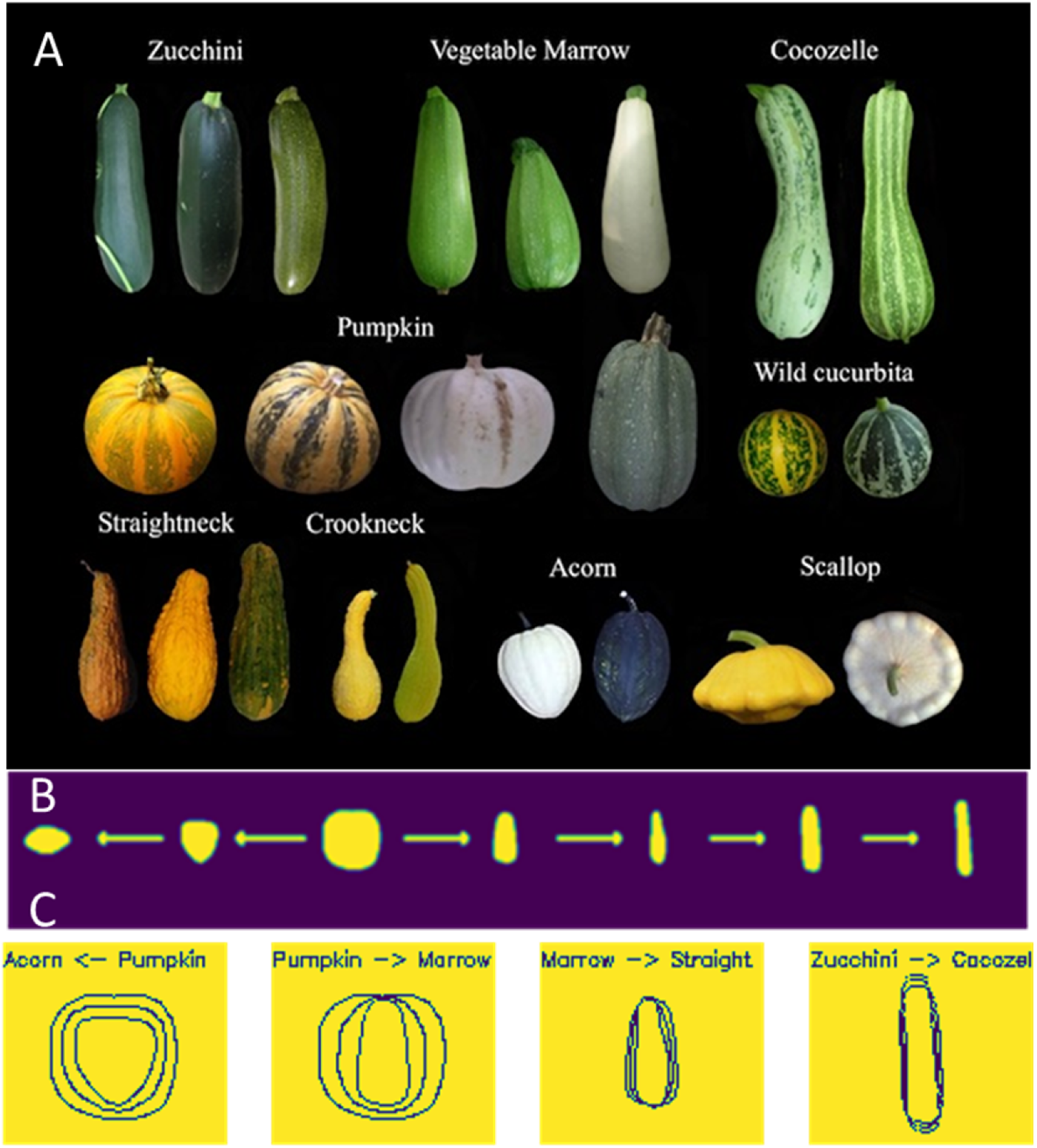
**A)** Variability found in C. pepo fruit shapes. B) Assumed evolutive pathways for shape simulation: scallop ← acorn ← pumpkin / wild gourd → marrow → straightneck → zucchini → cocozelle. **C)** Each panel shows contours of two observed shapes and an intermediate shape, illustrating how a continuous evolutive gradient corresponds to a given shape.

To generate *C. pepo* shapes along the putative evolutive gradient, we first sampled a random number from a uniform distribution *s* ~ U(−1, 1), where *s* = −1 defines an ‘acorn’ form; 0, a ‘pumpkin’, and 1 corresponds to ‘cocozelle’ (Figure 1B). Using the sampled *s* value, the two closest basic shapes were identified, and we defined a function that drew an intermediate shape between the nearest basic shapes, weighted by the proximity to each of the bounding contour (Figure 1C, see code in GitHub https://github.com/miguelperezenciso/dna2image/blob/main/dna2img.cucurbita.ipynb). The fruit corresponding to shape *s* was drawn in a 64 x 64 pixel image and noise was added to mimic rugosity of naturally observed fruits. This was done by adding an autoregressive noise to the contour (see code in GitHub). The decoder was trained using the ‘true’ *s* value and 100 random uncorrelated variables as input and the cucurbit shape images as output; 1,000 images were used for training and 100 for testing.

### Tomato shapes from experimental crosses

We used 353 tomato images from 129 crosses between 25 traditional varieties and 7 modern inbreds (Table S1). Traditional varieties were a subset of the TRADITOM project, which collected a wide sample of traditional tomato varieties from Southern Europe (Pons *et al*.2022; Blanca *et al*. 2022). Longitudinal cuts from about three fruits per parental or crossed line were photographed. Fruit images were segmented using a cluster algorithm (k=3) and contours were identified using a thresholding algorithm, as implemented in openCV. Contours were centered, cropped, and resized to 128 x 128 pixel binary images.

Modern inbred and traditional varieties were genotyped by sequence (GBS) previously (Blanca et al. 2022). Sixty eight segregating SNPs located within fruit shape candidate genes (Pons et al. 2022) were extracted. Hybrid offspring GBS genotypes were inferred from their parental genotypes. In addition, 48 biochemical, color and morphological metrics obtained with tomato analyzer had been obtained from each of the hybrid tomato fruits (Pons *et al*.2022) were also used for prediction. These metrics were not available for the 32 founder lines and were inferred with linear regression assuming additivity. This was done separately for each metric. The final network was trained using the 116 (68 + 48) ‘DNA’ measures as input for each of the accessions and the 353 tomato images as output. Input values were the same for images pertaining to the same accession.

### Shape prediction

We used a simple decoder architecture made-up of a first fully connected layer, followed by a reshaping layer and by three transposed convolutional layers (Figure 2). Code was implemented in keras and tensorflow (https://keras.io/, Abadi et al. 2015; Chollet 2015) and is inspired in autoencoder architectures (Brownlee 2019; Chollet 2021). The same decoder architecture was used for ellipse, cucurbita or tomato shape prediction, except that layer dimensions were adjusted according to image size (Figure 2). For ellipsoid 3D predictions, 3D transposed convolution layers were used instead of 2D transposed convolutions, but architecture was otherwise identical (see code in https://github.com/miguelperezenciso/dna2image).

**Figure 2:**
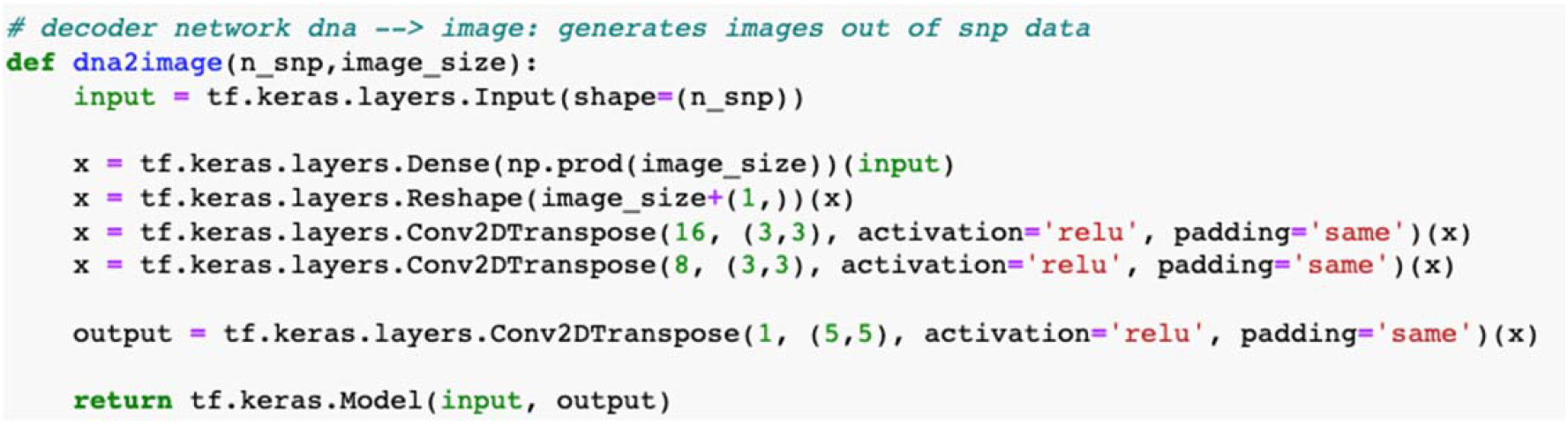
Keras code with the decoder used for image prediction. Function requires number of SNPs and output image size as input parameters.

### From phenotype to phenotype

Modern phenomics has sparked interest in ‘phenomic selection’, which consists in replacing genotyping by high throughput phenotyping to predict future offspring performance (Rincent *et al*. 2018; Cuevas *et al*. 2019; Robert *et al*. 2022). Here we considered two scenarios. In the first scenario we predicted 2D ellipses given two ‘parents’ ellipses. To do that, we first need to specify inheritance rules for images. Four arbitrary ‘image inheritance’ actions were defined:

- Additivity: the ‘offspring’ ellipse x and y coordinates are obtained by averaging coordinates of ‘parent’ ellipses.
- Dominance: for any pair of parent coordinates, the maximum of the two coordinates is selected as offspring coordinate.
- Imprinting: the offspring ellipse is identical to the first parental ellipse.
- Epistasis: the offspring ellipse is drawn by swapping the x and y coordinates of an ellipse intermediate between parents’ coordinates. That is, the epistatic offspring ellipse is a transposed additive ellipse.

We generated ~ 1,000 ellipse trios for each inheritance pattern to train the network. We trained the network for each inheritance pattern separately.

In the second, more realistic scenario, we used all combinations of male, female and offspring tomato images in a given cross from the previously described dataset. This resulted in a dataset of 2,325 tomato image trios. We utilized the same autoencoder architecture in both ellipse and tomato scenarios. Input consisted of two images that fed two separate CNN layers, one for each parental image, that were next concatenated (Figure 3).

**Figure 3:**
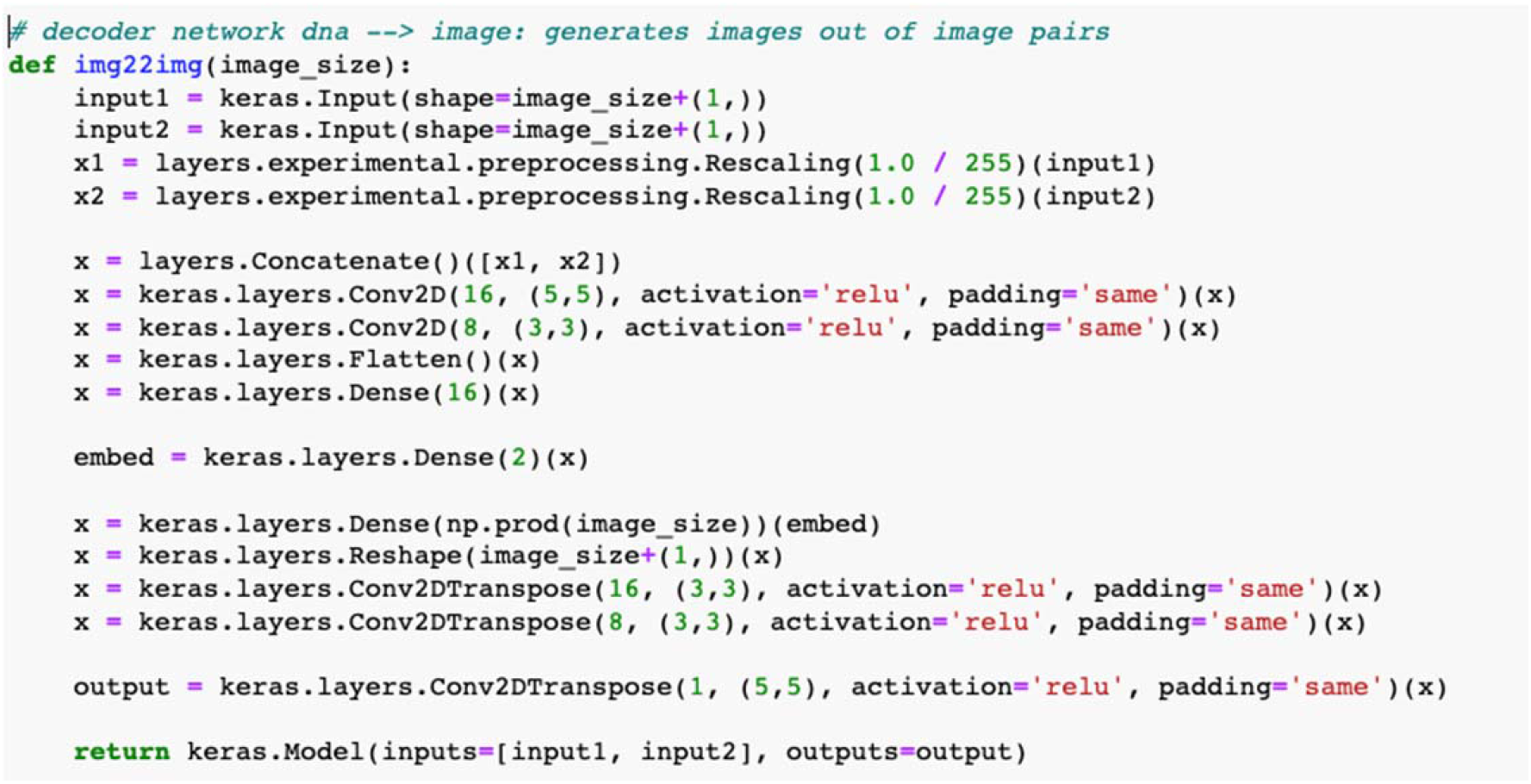
Keras code used for offspring image prediction based on parents’ images. It requires image size as input, which should be the same in input and output images. Size of embed vector can be fine-tuned for better performance.

## Results

### Shape prediction

We first show, as proof of concept in a toy example, that the simple decoder architecture in Figure 2 is able to learn and generate 2D and 3D simple forms from ‘genotype’ data. To train the decoder, we generated ~ 1000 2D ellipses and 3D ellipsoids with varying axis ratios and sizes (volumes) and the network was validated in 100 additional test images. Figure 4 show a sample of observed and predicted 2D ellipses, while results for 3D shapes are in Figure 5. In this latter case, sections across the three axes are shown for clarity since the 3D figure drawn with python package trisurf was not too accurate. Prediction is remarkably accurate also in the case of 3D shapes, especially when one considers the high dimensionality of the output image: 32 x 32 x 32 = 32,768 float numbers. Albeit in a simplistic scenario, we can see a naïve decoder is quite effective in predicting shapes conditional on text (DNA) information.

**Figure 4:**
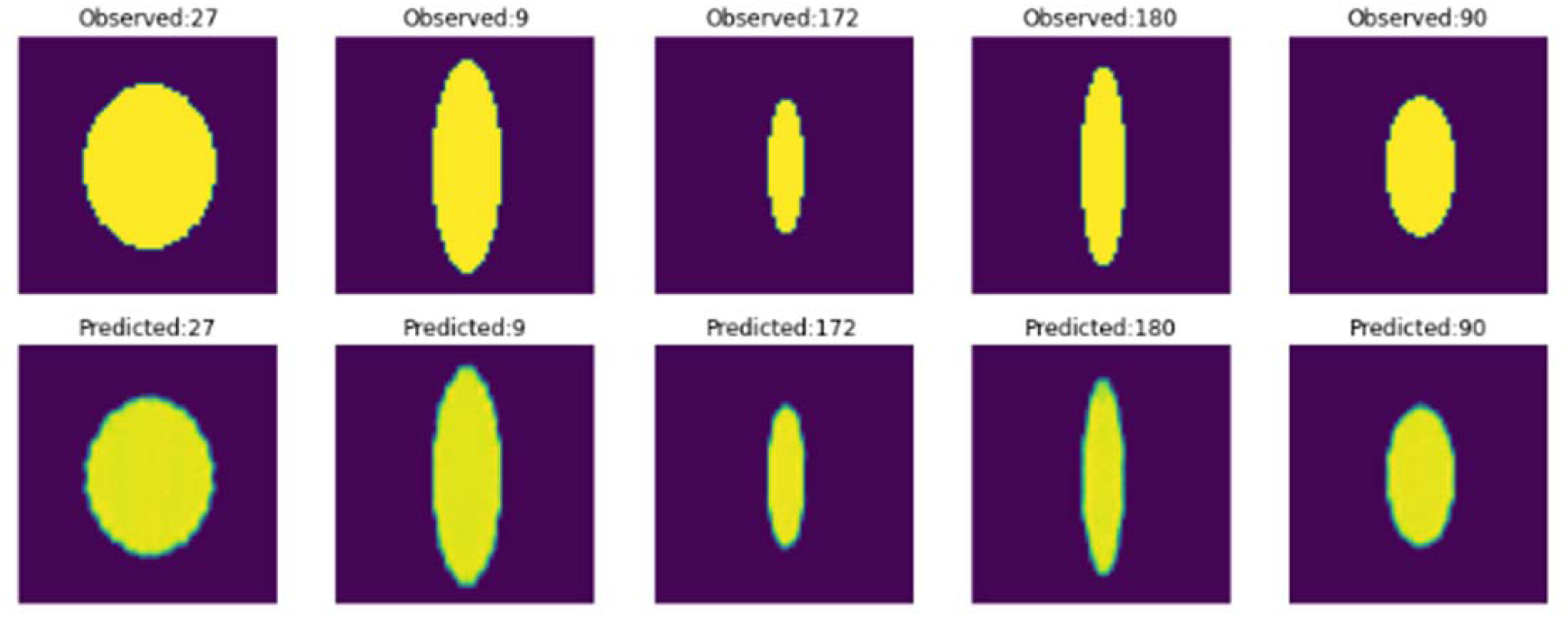
Top row: random sample of simulated ellipses; bottom row: predicted images using decoder in Figure 2.

**Figure 5:**
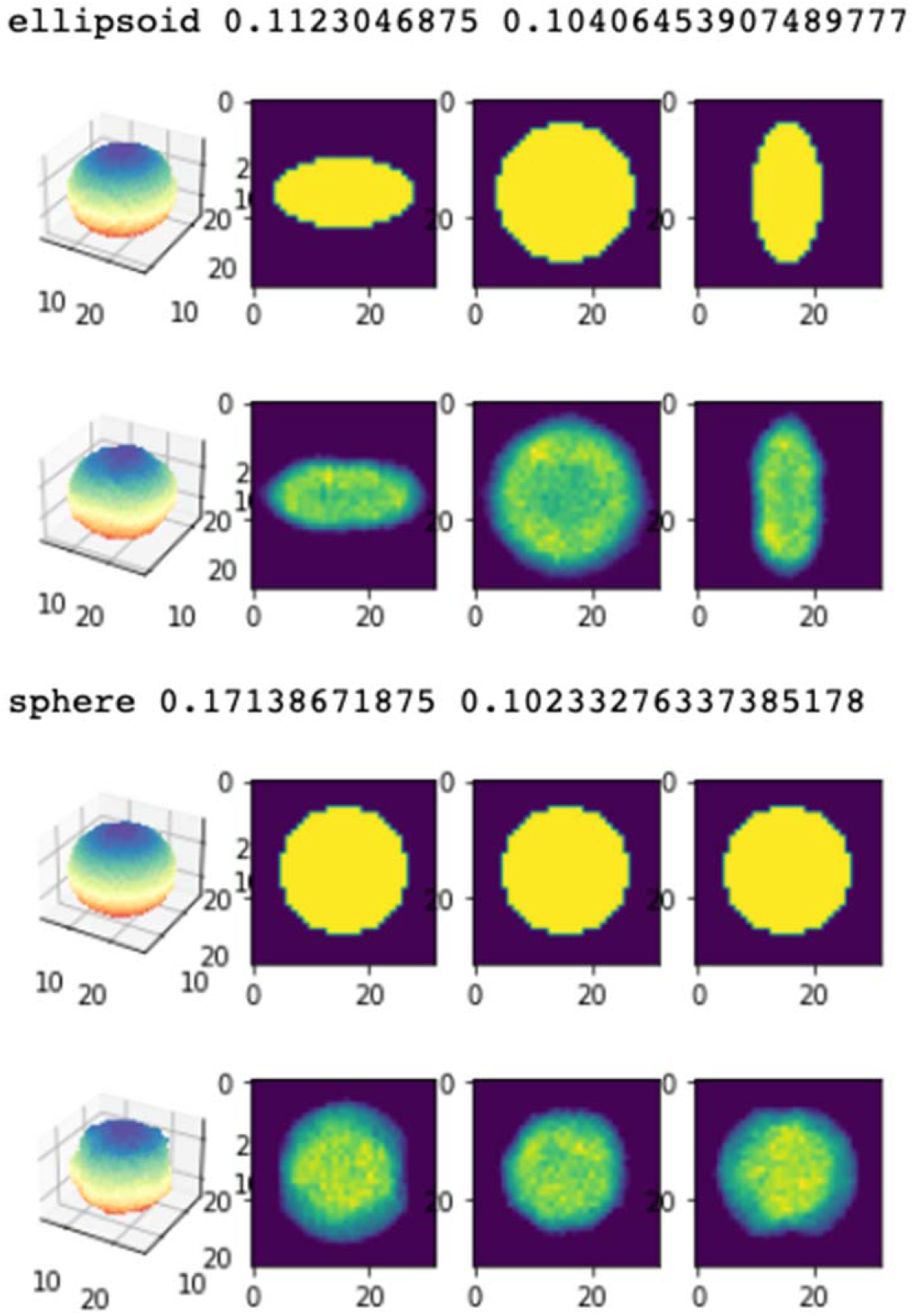
Generated (top rows) and predicted (bottom rows) of two 3D ellipsoids. The left column represents observed and predicted 3D representation, and the following columns are transversal cuts along the three axes.

To investigate whether the decoder network can be applied to more complex and realistic scenarios, we simulated cucurbit images from *C. pepo* as described in methods. We trained the same decoder as in the previous toy example using the shape causative locus *s* plus 100 random SNPs as input and the simulated cucurbit images as output. An example of five randomly predicted images is in Figure 6. Overall, prediction was quite reasonable, and predicted shapes can be easily recognized. Note the ‘rugosity’ induced by the autoregressive model, which is also reproduced in the prediction. We found the maximization algorithm can have a large influence on results. RMSprop performed best, whereas Adam failed often and Adagrad did not seem to work.

**Figure 6:**
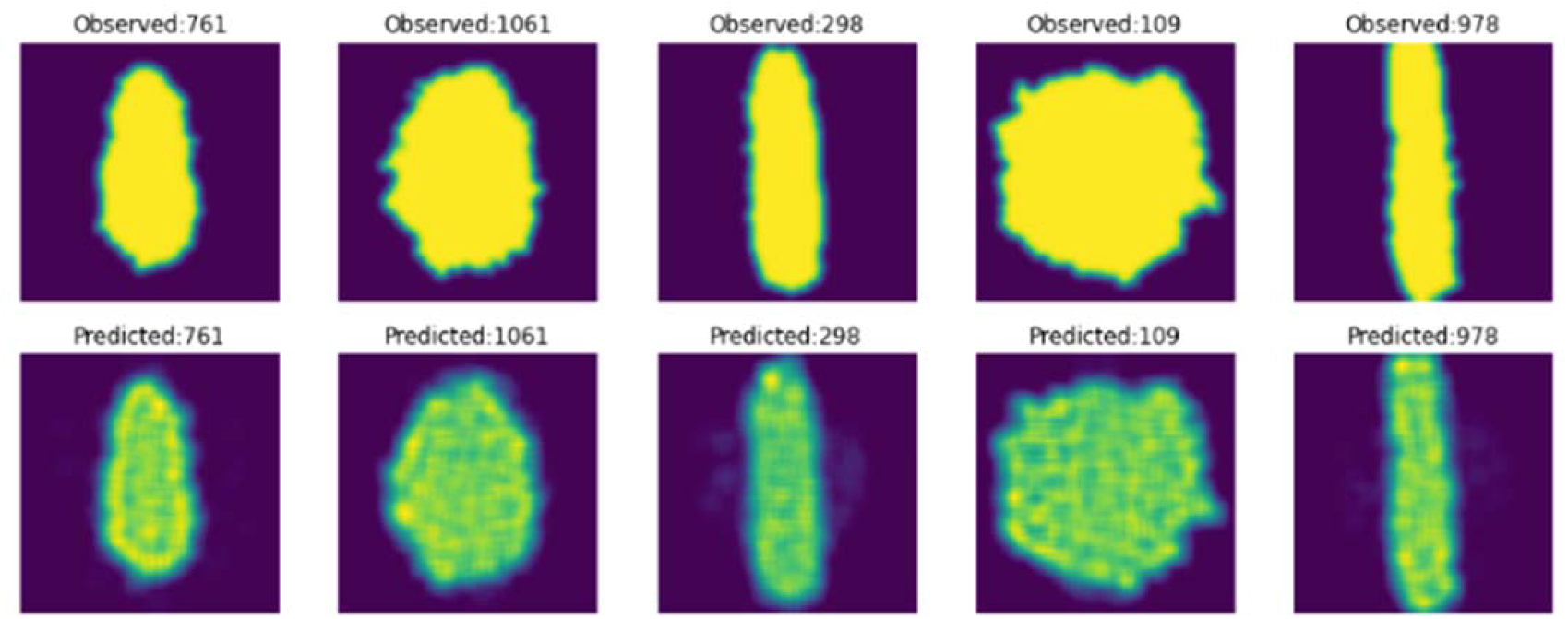
Top row: sample of simulated cucurbit images including autoregressive noise; bottom row: predicted images using decoder in Figure 2.

Prediction of a random set of tomato shapes based on the 116 metrics is shown in Figure 7. Predictions were very good overall, except of hybrids involving TR_MO_004 (Figure 7, sample 1). This traditional variety belongs to the horticultural group “Coeur de Boeuf”, which fruits are big with irregular shapes.

**Figure 7:**
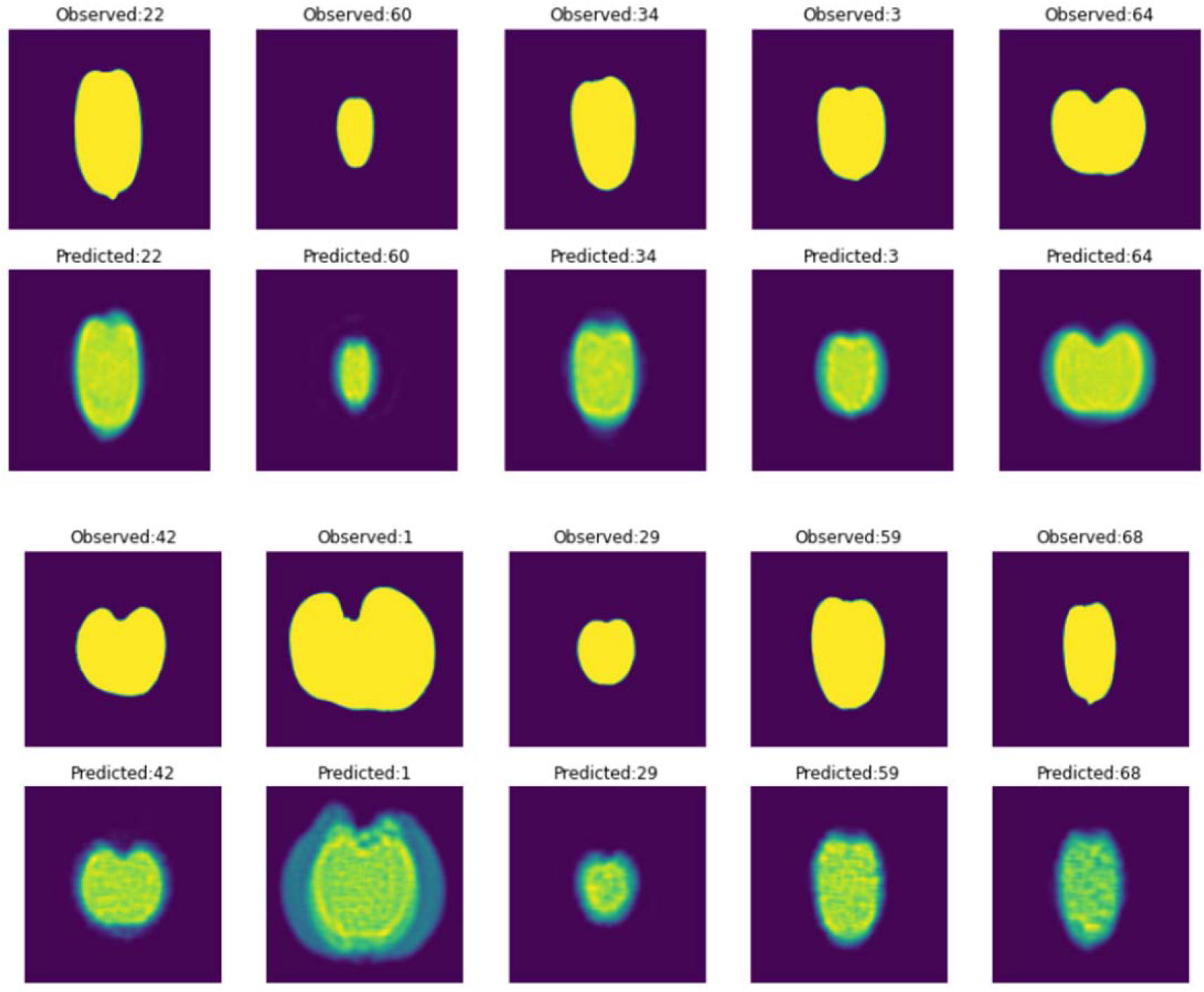
Sample of observed tomato images (first and third rows) and the corresponding predicted images using decoder in Figure 2.

### From phenotype to phenotype

Can we bypass genotype information altogether? If shapes are highly heritable, the network could learn inheritance patterns and predict offspring shape directly from parents’ forms, without resorting to genotypes. Figure 8 shows examples of the four image ‘inheritance’ behaviors defined: ‘additivity’, ‘dominance’, ‘imprinting’ and ‘epistasis’. We observe that predictions were reasonably accurate for additivity and epistasis but were worse for dominance and, especially, for imprinting. It seems the network can accurately find additive and non-linear patterns but is less adapted to predictions where the order of inputs is relevant. We conjecture then that recurrent neural networks (RNNs, e.g., Hill et al. 2018) could be better suited to this problem.

**Figure 8:**
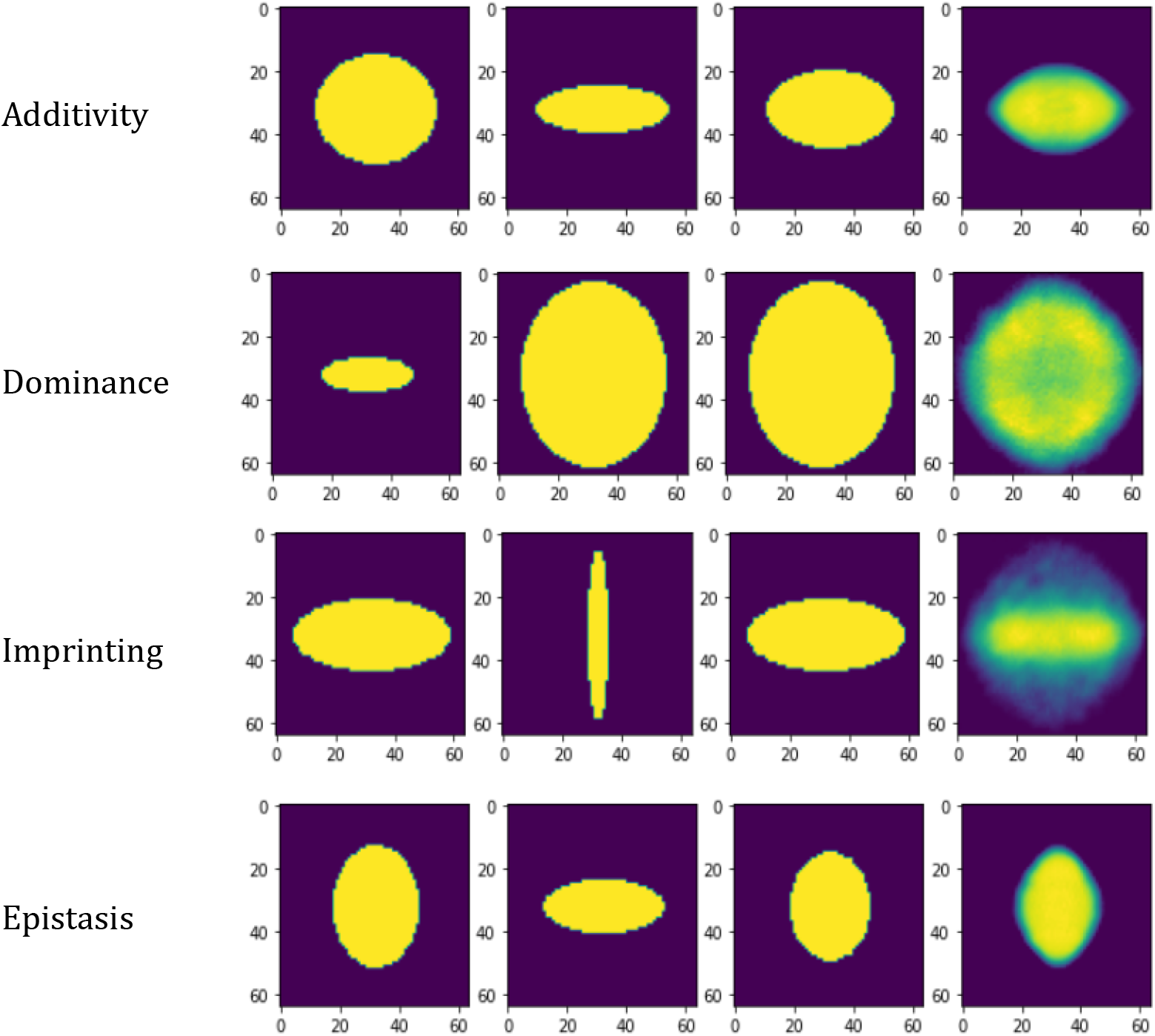
Examples of the four arbitrary image inheritance patterns defined: ‘additivity’, ‘dominance’, ‘imprinting’ and ‘epistasis’. Columns show ‘paternal’, ‘maternal’, ‘offspring’ and predicted images.

In the second example, we used the images from crosses between traditional and modern inbred tomato lines described. Predictions (Figure 9) were remarkably accurate overall, proving fruit appearances can be predicted from ancestor images. It also suggests that the predominant action seems to be additive.

**Figure 9:**
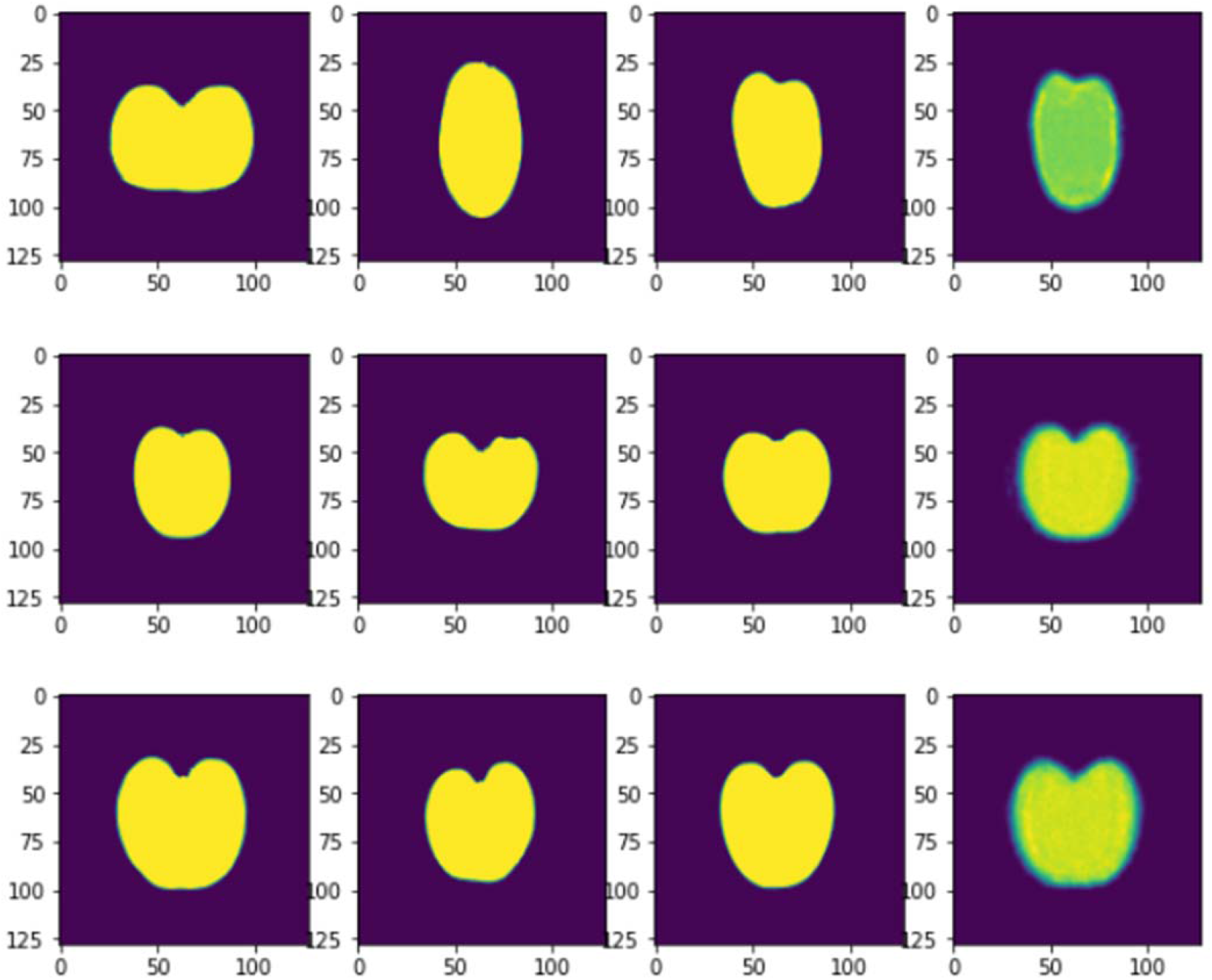
Observed tomato trios in three random crosses and predicted offspring based on network in Figure 4. Columns are paternal, maternal, offspring and predicted offspring images. Images’ size is 124 x 124 pixels.

## Discussion

Being able to predict highly dimensional objects such as appearance can revolutionize breeding by merging genome and phenome information in a coherent framework. Here we present a proof of concept that this is possible, even using very simple network architectures. We show that 2D, but also 3D, shapes can be accurately predicted and generated.

The problem posed here is similar to the ‘text-to-image’ challenge, where algorithms are trained to generate images from figure captions. Some works have recently reported highly accurate results (Ramesh *et al.*; Radford *et al*. 2021) and we foresee that ‘dna-to-image’ should follow. There are some differences between text and DNA that require specific developments though. First, text is divided in a finite, relatively small number of items (words) which relationships can be inferred by automatically parsing large available databases. DNA sequence can be split into coding / noncoding, introns / exons but cannot be assimilated to ‘words’ with specific meanings. DNA or marker data are not segmented; their relationships are much more intricate than those in words from human languages and are unknown to a large extent. For instance, most discovered causative mutations that affect shape are located outside coding regions (Wu *et al*. 2018; Martínez-Martínez *et al*. 2022). Second, large corpuses of images and figure captions are available for training text-to-image problems; these datasets are not readily available for fruits or other agricultural scenarios. Finally, texts used to generate images are short and simple; algorithms usually fail and generate unpredictable results if input text is slightly changed. In the case of DNA, the number of differences between strain or individual genotypes is very large; we still do not know how dna-to-image algorithms will cope with this issue.

Text-to-image methods rely on text encoding, also called ‘embedding’, i.e., in finding an optimum numeric representation of text elements in a reduced n-dimensional space. DNA encoding is to be critical in dna-to-image problems as well. Previous research on DNA encoding has utilized small DNA sequences, e.g., taking exons as ‘words’ (Zou *et al*. 2019; Ji *et al*. 2021). However, this cannot be applied to generic marker data or complete sequence. We hypothesize that standard dimension reduction techniques, such as classical principal component analysis (PCA), can be a useful alternative especially when shape is controlled by numerous loci of small effect.

For simulation purposes of cucurbit shapes, we assumed an underlying continuous gradient that results in a continuous morphological variation (Figure 1C). We assumed this for computational and illustrative purposes, although we reckon there is no clear biological evidence on this hypothesis. Modern cultivars adopt discrete shapes and intermediate shapes are rarely observed. However, traditional unimproved varieties and their crosses do show a number of intermediate features (Montero-Pau *et al*. 2017).

Numerous genes that influence shape have been discovered (Monforte *et al*. 2014; Grumet and Colle 2016; Snouffer *et al*. 2020). These genes act in concerted action during development (Wu *et al*. 2018). Note the method proposed here does not require causative loci to be identified, as prediction methods rely on linkage disequilibrium between causative and genotyped markers. Nevertheless, known causative polymorphisms could be given larger weights than the rest of SNPs. There are several approaches that can be used to achieve this. One option is the ‘attention’ mechanism, which is used to underline words of particular relevance in text analysis (Vaswani *et al.*). Another possibility is to define a specific input layer for causative mutations and merging with the rest of SNPs in a separate layer. This is straightforward with standard software such as Keras (Chollet 2015).

Further work is warranted to overcome limitations of this work and continue this area of research. First of all, appropriate datasets of large size in 2D and 3D must be generated. In fact, one of the limiting steps for this methodology to be applied is the lack of datasets of enough size containing high density genotypes and good quality images. The simplest scenario should be fruits, as is the TRADITOM initiative in tomato (Pons *et al*. 2022; Blanca *et al*. 2022), but many other applications can be envisaged: animal conformation (e.g., dairy bull catalogs, dog breeds), whole plant appearance, leaf and root morphology, color patterns,… Second, more complex network architectures inspired in current text-to-image algorithms must be adapted to the dna-to-image scenario. Finally, generative models, such as conditional generative adversarial networks (CGANs; Goodfellow et al. 2014; Mirza and Osindero 2014), conditional on DNA information, could be used to produce images of high quality. On top of that, new tools for dealing with 3D objects are needed.

In summary, we have shown that very simple networks can be successfully trained in small datasets to accurately predict fruit images. Although much work remains to be done, this research opens new possibilities in the area of prediction of complex traits.

## Data availability statement

All data and code are available at https://github.com/miguelperezenciso/dna2image.

## Acknowledgments

Authors acknowledge the technical support of Gorka Perpiña and Eva María Pérez. Modern inbreds and their hybrids with traditional tomato varieties were provided by Meridiem Seeds (https://meridiemseeds.com/, Murcia, Spain).

## Funding

Work funded by Ministry of Science and Innovation-State Research Agency (AEI, Spain, 10.13039/501100011033) grant numbers PID2019-108829RB-I00 and CEX2019-000902-S, European Commission H2020 research and innovation program through TRADITOM grant agreement No.634561, and HARNESSTOM grant agreement No.101000716, and PROMETEO projects 2017/078 and 2021/072 to promote excellence groups by the Conselleria d’Educació, Investigació, Cultura i Esports (Generalitat Valenciana, Spain) and by the CERCA Programme / Generalitat de Catalunya (Spain).

## Author contributions

MPE and LMZ conceived computational research and developed code; AG, BP and AJM conceived and discussed empirical research; MPE, CP and SS performed research. MPE and AJM wrote the draft with help from rest of authors.

---

**Table S1:** Parental tomato lines.

| Code | Type | Fruit type |
| --- | --- | --- |
| MS_1 | Modern Inbred | Salad tomato |
| MS_2 | Modern Inbred | Salad tomato |
| MS_3 | Modern Inbred | Salad tomato |
| MS_4 | Modern Inbred | Long processing |
| MS_5 | Modern Inbred | Cocktail round |
| MS_6 | Modern Inbred | Cherry round |
| MS_7 | Modern Inbred | Cherry round |
| TR_TH_001 | Traditional | round |
| TR_TH_002 | Traditional | round |
| TR_TH_003 | Traditional | flattened |
| TR_CA_001 | Traditional | obovoid |
| TR_CA_002 | Traditional | flat |
| TR_VA_001 | Traditional | flat |
| TR_VA_002 | Traditional | oxheart |
| TR_VA_003 | Traditional | round |
| TR_MO_001 | Traditional | flat |
| TR_MO_002 | Traditional | round |
| TR_MO_003 | Traditional | round |
| TR_MO_004 | Traditional | oxheart |
| TR_VI_001 | Traditional | Long |
| TR_VI_002 | Traditional | round |
| TR_VI_003 | Traditional | round |
| TR_VI_004 | Traditional | ellipsoid |
| TR_VI_005 | Traditional | obovoid |
| TR_VI_006 | Traditional | rectangular |
| TR_PO_001 | Traditional | ellipsoid |
| TR_PO_002 | Traditional | long |
| TR_PO_003 | Traditional | ellipsoid |
| TR_PO_004 | Traditional | ellipsoid |
| TR_IS_001 | Traditional | long |
| TR_IS_002 | Traditional | obovoid |
| TR_IS_003 | Traditional | round |

MS\_1 to MS\_7 correspond to modern inbred lines provided by Meridiem Seeds. Codes for the traditional varieties are according TRADITOM project (Blanca et al. 2022).

